# Human iPSC-derived mesodermal progenitor cells preserve their vasculogenesis potential after extrusion and form hierarchically organized blood vessels

**DOI:** 10.1101/2021.05.09.443303

**Authors:** Leyla Dogan, Ruben Scheuring, Nicole Wagner, Yuichiro Ueda, Philipp Wörsdörfer, Jürgen Groll, Süleyman Ergün

## Abstract

Post-fabrication formation of a proper vasculature remains an unresolved challenge in bioprinting. Established strategies focus on the supply of the fabricated structure with nutrients and oxygen and either rely on the mere formation of a channel system using fugitive inks, or additionally use mature endothelial cells and/or peri-endothelial cells such as smooth muscle cells for the formation of blood vessels *in vitro*. Functional vessels, however, exhibit a hierarchical organization and multilayered wall structure that is important for their function. Human induced pluripotent stem cell-derived mesodermal progenitor cells (hiMPCs) have been shown to possess the capacity to form blood vessels *in vitro*, but have so far not been assessed for their applicability in bioprinting processes. Here, we demonstrate that hiMPCs, after formulation into an alginate / collagen type 1 bioink and subsequent extrusion, retain their ability to give rise to the formation of complex vessels that display a hierarchical network in a process that mimicks the embryonic steps of vessel formation by vasculogenesis. Histological evaluations at different time points of extrusion revealed initial formation of spheres, followed by lumen formation and further structural maturation as evidenced by building a multilayered vessel wall and a vascular network. These findings are supported by immunostainings for endothelial and peri-endothelial cell markers as well as electron microscopic analyses at the ultrastructural level. Moreover, capillary-like vessel structures deposited a basement membrane-like matrix structure at the basal side between the vessel wall and the alginate-collagen matrix. These results evidence the applicability and great potential of hiMPCs for the bioprinting of vascular structures mimicking the basic morphogenetic steps of *de novo* vessel formation during embryogenesis.

## 1. Introduction

Vascular disorders are the main reason of cardiovascular diseases, e.g. coronary artery disease, that are leading death cause worldwide. The most used clinic approach to overcome vascular failure is the transplantation of autologous vessel pieces as vascular graft, e.g. using internal thoracic artery or saphenous vein for coronary bypass surgery. Not only for treatment of such vascular diseases but also for tissue regeneration and replacement the functional vascularization is an indispensable component for achieving suited internal tissue structure in biofabricated cell-material composites. Hence, there is a huge need for approaches enabling the engineering of blood vessels regardless with which technology. First steps of creating artificial blood vessels were undertaken by tissue engineering approaches, e.g. cell seeding into the decellularized matrix [1], annular mode casting [2], seeding endothelial cells into tubular structures that was created using collagen or collagen with some other matrix components like elastin [3-5]. Further attempts were made using biomaterials that were constructed as basic vascular structures and again seeded with multiple cell types [6, 7]. While these approaches resulted in tubular conduits the spatial distribution of the used cell types as well as the scaffold architecture remained incomplete. More recently, bilayered small vessels were bioprinted using HUVECs (human umbilical vein endothelial cells) and smooth muscle cells (SMCs) [8].

Bioprinting is a novel promising option to create artificial blood vessels. While great progress has been achieved in biofabrication of some functional tissues in the past decade, bioprinting of an appropriate vascular system beyond mere generation of channel structures is still a big challenge. A scaffold-free tubular tissue was created using 3D-bioprinting of multicellular spheroids that were composed of human umbilical vein endothelial cells (HUVEC), aortic smooth muscle cells SMCs) and human dermal fibroblasts [9]. The functionality of these structures was tested *in vivo* after implantation into the rat abdominal aorta [9]. Few years later, the same group demonstrated the formation of tubular structures that were lined by endothelial cells using bioprinting of a cellular mix composed of endothelial cells, dermal fibroblast and iPS-derived cardiomyocytes [10]. Recently, 3D bioprinting of a bilayered vessel-like tubular structure was demonstrated using the extrusion-based bioprinting of HUVECs and SMCs into Gelatin Methacryloyl (GelMA) as hydrogel with varying porosity (Lei et al., Biofabrication, 2020). This approach resulted in spatial separation of the printed cells in tubular structures that exhibited an inner layer containing HUVECs and an outer layer displaying SMCs.

While the aforementioned approaches delivered vascular-like 3D tubular structures, there are still major missing structural properties when compared to the vascular system *in situ*. The mature vascular system is composed of a complex network of various sized blood vessels that are organized in a hierarchical order: large or mid-sized arteries and veins are connected to each other by microvessels such as arterioles, capillaries and venules. Except for capillaries, all other types of blood vessels display a three-layered wall with the innermost intima layer that contains ECs and followed by the media layer which is constructed by SMCs that provide the contractility of the vessel wall and the outermost adventitial layer [11, 12]. Capillaries also contain an intima that is lined by ECs and enwrapped by a discontinuous layer of pericytes form the outside. While both vascular intima and media are constructed by mature cells, the outermost vascular adventitia has been identified as a niche for vascular stem and progenitor cells (VW-SCs) that can deliver all vascular cells and some types of blood cells upon activation, e.g. by angiogenic stimuli [11, 13-15]. Considering this complexity of the *in situ* vascular system, it will probably not suffice when endothelial tubes alone or with some covering SMCs are printed.

In this current manuscript, we pursue an alternative approach for creating a complex and hierarchically organized 3D vascular network that resembles the vascular system as closely as possible *in situ*. Therefore, we used human iPSC-derived mesodermal progenitor cells (hiMPCs), that were shown to deliver both ECs and pericytes [16], instead of often used mature ECs, SMCs or fibroblasts, and formulated them in alginate+collagen type I hydrogels. We then extruded these bioinks into molds to evaluate whether a *de novo* embryonic development of a vascular system can be achieved. We demonstrate here, that hiMPCs survive the extrusion into the alginate alone or alginate+collagen type I hydrogel and subsequently pass through a series of morphogenetic events under culture conditions starting mostly at culture days 5-6 after extrusion. More intriguingly, these morphogenetic events begin with the formation of cell spheroids from which the nascent endothelial tubes form by self-assembly. These primitive vessel-like structures undergo several steps of further maturation and stabilization by assembling of additional vascular mural cells such as pericytes or SMCs. Moreover, these vessel-like structures create a network that is composed of vessels showing different diameter and varying wall structure: capillary-like vessels displaying ECs, pericytes and a basal lamina as well as three-layered vessel-like channels displaying an intima, media and adventitia as evidenced by electron microscopic analyses. Consistent with these data, immunofluorescence analyses confirmed the luminal presence of CD31^+^ ECs, that were covered by NG2^+^ (pericyte marker) or a-SMA^+^ (smooth muscle cell marker) cells from the outside. Moreover, CD34^+^ cells were found in the outermost layer of these vessel-like structures similar to the blood vessels *in situ* that harbor CD34^+^ cells in their adventitial layer [17-19].

Taken together, this pioneering study reveals the suitability of hiMPCs for extrusion based bioprinting and elaborates suitable biomaterial properties that allow hiMPCs to unleash their vasculogenic potential closely mimicking basic steps of embryonic *de novo* vascular development. We show that hiMPCs have the capacity to deliver all vascular cell types which self-assemble into multi-layered vessel-like structures that are capable of forming a hierarchically organized vascular network.

## 2. Materials and methods

### 2.1 Cell Culture

Human iPSCs (hiPSCs) and bioinks containing hiPSC-derived mesodermal cells (hiMPCs)[16] were used for the extrusion-based dispensing into silicone molds. The hiPSCs were generated from commercially available normal human dermal fibroblasts (juvenile NHDF, C-12300, Promocell, Heidelberg, Germany) by reprogramming, using either the hSTEMCCA-lentiviral construct [20, 21] or a Sendai virus reprogramming kit (Cyto Tune 2.0 Sendai reprogramming vectors, Ref#A16517; Lot#A16517, Invitrogen, Carlsbad, CA). hiPSCs were cultured on hESC-qualified Matrigel^®^ (Corning, New York, NY, USA)-coated culture plates in StemMACS iPS Brew medium (Myltenyi Biotec, Bergisch Gladbach, Germany). The culturing media was changed daily. At ∼80-85% confluency, cells were dissociated with StemPro® Accutase® (Thermo Fisher Scientific, Waltham, MA, USA) for 5 min at 37°C and triturated to obtain a single cell suspension. hiPSCs were plated at a cell density of 2×10^4^/cm^2^ in StemMACS medium, supplemented with 10 µM rho-associated protein kinase inhibitor Y-27632 (Miltenyi Biotec) for the first 24 h after passaging. Basic characterization of the hiPSC line has been done by checking the expression of the pluripotency markers Oct4 and Sox2 (Fig.1 A1-A3) using immunofluorescence staining (IF). Subsequently, the hiPSCs were converted to hiMPCs under directed differentiation conditions as recently published [16]. In brief, confluent hiPSCs were dissociated into a single cell suspension using Accutase and were counted. For mesodermal induction, 3.65×10^4^ hiPSCs per cm^2^ were seeded on Matrigel-coated plates and cultured for one day in StemMACS iPS-Brew medium with 10µM protein kinase inhibitor (Y-27632). Subsequently, StemMACS iPS-Brew medium was replaced by mesodermal induction medium (Advanced DMEM/ F12, 0.2mM L-Glutamine, 60µg/µl Vitamin C, 10µM CHIR 99021, 25 ng/ml BMP4) and the cells were cultured for further 3 days at 37°C, 5% CO_2_. Culture medium was changed daily. After 3 days of culturing in the mesodermal induction media, the cells were dissociated using Accutase and collected for extrusion-based dispensing of bioinks containing hiMPCs into the molds.

**Figure 1:**
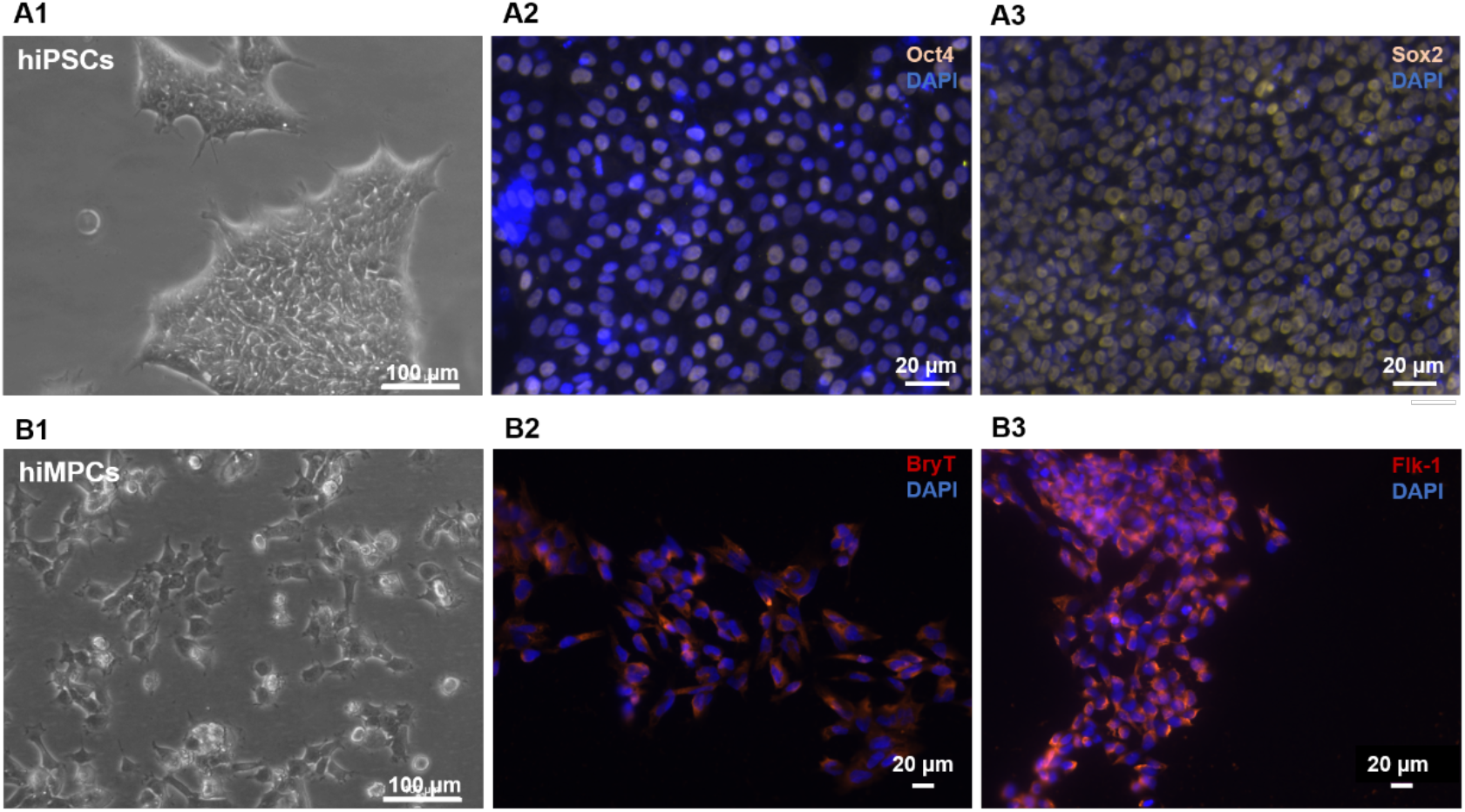
**A1** and **B1**. Phase contrast microscopic images of hiPSCs and hiMPCs respectively, cultured in standard 2D culture according to protocol described in materials and methods section. **A2**. and **A3**. Immunofluorecence staining of hiPSCs for Oct4 and Sox2. (B2, B3). Immunofluorescence staining of hiMPCs for BryT and Flk1.

After extrusion, the cell-loaded discs were cultured with specified medium at 37°C, 5% CO_2_. To induce the differentiation of the extruded hiMPCs into vascular cells e.g. endothelial cells (ECs), vascular differentiation medium containing VEGF was used (Advanced DMEM/F12, FCS, Penstrep, 10 µM ROCKinhibitor (Y-27632), 0.2 mM L-Glutamine, 60 µg/ml Vit.C, 50 ng/ml VEGF).

### 2.2 Hydrogel preparation

Sodium alginate (Sigma Aldrich, PH132S2) and collagen type I (Millipore, E2729) were used for bioink formulation. To prepare 2% (wt/vol) alginate solution, sodium alginate powder was weighed, placed in a glass vial and sterilized for 1h under UV light. Then the sterilized powder was dissolved in PBS (Sigma-Aldrich) solution by stirring over night at 37 °C. Collagen type 1 solution was dispersed in advanced DMEM, then 5N NaOH (Sigma, USA) was used to adjust the pH=7. All bioinks were stored in sterile conditions at proper temperature. Prior to extrusion, hiMPCs were immediately dispersed within the collagen type 1(0.015 % w/v) and subsequently, this dispersion of cells was mixed with alginate carefully. Then this cell-loaded hydrogel mixture was loaded into syringes and subsequently extruded into the silicon molds to form spherical discs with 80 microliter cell-loaded gel volume. In terms of using of alginate+collagen type I hydrogel, cell suspension was prepared at 4°C using the appropriate (0.015 % w/v) amount of collagen type I. The prepared collagen type1+cell mixture was added to the alginate carefully. A homogeneous gel mixture was prepared by controlled up and down pipetting without creating bubbles and the bioink was immediately extruded under the same extrusion conditions like in pure alginate case. Over culturing period, it was observed that the extruded hiMPCs cells were viable and functional within the hydrogel mixture. The cellular behavior was followed by cell viability assays and the viability ratios were assessed at three incremental time points: 1, 3, and 7 days.

### 2.3 Immunocytochemistry analysis

To characterize the cells before printing, the cells (cultured in 2D) were fixed with 4% paraformaldehyde (AppliChem, Darmstadt, Germany) in PBS for 15 min at room temperature (RT), then blocked with blocking buffer solution 5% NGS (Normal Goat Serum) for 1h at the same temperature, in the presence of 0.1%Triton X-100 (Sigma–Aldrich) to permeabilize the cell membrane for detection of intracellular markers, or without Triton-X for detection of cell surface markers. Thereafter, the cells were incubated with primary antibodies in 2%BSA (Bovine Serum Albumin) +1% NGS overnight at 4°C. For immunocytochemical characterization of hiPSCs and hiMPCs following primary antibodies were used: Oct4 (Santa Cruz Biotechnology,sc-6279), Sox2 (R&D Systems, MAB2018), Brachyury (T) (R&D Systems, AF2085), Flk1 (Santa Cruz Biotechnology, sc-48161). Cells were incubated with primary antibodies overnight at 4 °C. The incubated samples were rinsed 3 times with PBS, and exposed to appropriate fluorophore-conjugated secondary antibodies that were diluted in the PBS solution for 1h at RT. Secondary Cy2-, Cy3- or Cy5-labelled antibodies were used to visualize primary antibodies. The immunostained samples were then studied microscopically and used for capturing of pictures by the Biorevo fluorescence microscope (Keyence, Osaka, Japan). The analyses revealed that the hiMPCs express the mesodermal progenitor markers BryT and Flk1 (Fig.1 B1-B3).

For conventional histological analyses, the disc shaped scaffolds were removed from the culture and washed in PBS 3x for 5min. The scaffolds were fixed in 4% PFA that was freshly prepared in PBS for 24h at 4°C and then washed again 3x with PBS for 30mins to remove residual PFA. The fixed samples were stored submerged in %70 ethanol until embedding into the paraffine. Afterward, paraffin sections were prepared with required thickness (5-12 µm) and were then deparaffinized, rehydrated and stained with hematoxylin and eosin (H&E) or picrosirous red to visualize the collagenous matrix. For immunofluorescence analyses, antigens were unmasked using Sodium Citrate buffer (10 mM, pH6). For immunostaining, the sections were exposed to antibodies for following markers: CD34 (Miltenyi, 130-105-830), CD31 (DAKO, M0823), NG2 (Merck-Millipore, AB5320) (Abcam, AB5694). These markers are used at the most for detection and characterization of vascular wall cells. The stained sections were studied microscopically and used for capturing of pictures by the Biorevo fluorescence microscope (Keyence, Osaka, Japan).

### 2.4 Transmission electron microscopy

Extruded discs were fixed in fixation solution (0.15M cacodylate buffer pH 7.4 containing 2.5% glutaraldehyde, 2% formaldehyde with 2mM calcium chloride) on ice for 2 hours and washed 5 x 3 minutes in cold 0.15M cacodylate buffer (50mM cacodylate, 50mM KCl, 2.5mM MgCl_2_, 2mM CaCl_2_ pH 7.4) on ice. Subsequently, specimens were incubated on ice in a reduced osmium solution containing 2% osmium tetroxide, 1.5% potassium ferrocyanide, 2 mM CaCl_2_ in 0.15 mM sodium cacodylate buffer (pH 7.4). Specimens were washed with dH_2_O at room temperature (RT) 5 x 5 minutes followed by incubation in 1% thiocarbohydrazide (TCH) solution for 25 minutes at RT. Specimens were washed with ddH_2_O at RT, 5 x 5 minutes each and incubated in 2% osmium tetroxide in ddH_2_0 for 30 minutes at RT. Afterwards, specimens were incubated in aqueous UAR-EMS (4%, Uranyl Acetate Replacement stain, Electron Microscopy Sciences, Hatfield, USA) and stored at 4°C overnight. The next day, specimens were washed 3 x 3 minutes in ddH_2_O at RT. Prior incubation with lead aspartate solution, specimens were washed 2 x 3 minutes in ddH_2_O at 60°C and subjected to *en bloc* Walton’s lead aspartate staining [22] and placed in a 60°C oven for 30 minutes. Specimens were washed 5 x 5 minutes with ddH_2_O at RT and dehydrated using ice-cold solutions of freshly prepared 30%, 50%, 70%, 90%, 100%, 100% ethanol (anhydrous), 100% aceton (anhydrous) for 10 minutes each, then placed in anhydrous ice-cold acetone and left at RT for 10 minutes. Specimens were placed in 100% aceton at RT for 10 minutes. During this time, Epon812 was prepared. The resin was mixed thoroughly and samples were placed into 25% Epon:acetone for 2 hours, then into 50% Epon:acetone for 2 hours and 75% Epon:acetone for 2 hours. Specimens were placed in 100% Epon overnight. The next day, Epon was replaced with fresh Epon for 2 hours and specimens were placed in beem capsules and incubated in a 60°C oven for 48 hours for resin polymerization. For ultrathin sections, 70nm thick ultrathin sections were cut with an ultramicrotome (Ultracut E, Reichert Jung, Germany) and collected on copper or nickel grids and finally analyzed with a LEO AB 912 transmission electron microscope (Carl Zeiss Microscopy GmbH, Germany).

### 2.5 Serial block face electron microscopy (SBF-SEM)

For SBF-SEM, specimens were mounted on on aluminium pins (Gatan Inc). The blocks were precision trimmed with a glass knife to expose the spheroids. Silver paint (Gatan Inc) was used to coat the edges of the tissue block to reduce charging during imaging using BSE mode using the variable pressure mode. Images were acquired using a scanning electron microscope (Sigma300VP; Carl Zeiss Microscopy GmbH, Germany) equipped with an automated ultramicrotome inside the vacuum chamber (3View; Gatan Inc). The ultramicrotome cut successive sections at a thickness of 50 nm. After each section, the sampleblock-face was scanned. The microscope, the stage and the ultramicrotome were controlled using DigitalMicrograph software GMS3 (Gatan Inc). The SEM was operated in variable pressure mode (VP-Mode) The sample was scanned in VP-Mode with a chamber pressure of 15 Pa and a landing energy of 3,5 kV. Imaging in VP-Mode reduced artifacts arising from sample charging due to high amounts of epon between the spheroids.

#### 2.5.1 Image processing and segmentation

Image alignment was performed using Digital Micrograph (Gatan Inc). Segmentation was performed using the TrakEM2 plug-in [23] of the open source image processing framework Fiji [24].

#### 2.5.2 Data visualization

3D rendering model and movie (Supplementary Movie) were created using the open-source platform tomviz (https://tomviz.org/)

### 2.5 Cell Viability

Live/dead cell staining was performed for testing the cell viability using Calcein AM (2µM, C1430 Life technologies) and Ethidium Homodimer-1, EthD-1, (E1169, 4µM, Life technologies) in accordance with the manufacturer’s instruction. Briefly, three independently extruded discs were washed 3x by PBS for 5mins to remove the residual cell culture medium. Enough volume of freshly prepared staining solutions was added to cover the discs and then the discs were incubated at 37°C for 30mins. Afterwards, staining solutions were removed and replaced by PBS. The constructs were studied microscopically using a Leica PL microscope that is equipped with digital camera allowing to capture pictures. In addition, to count the cells three independent scaffolds were dissolved by 2mL of 20mM EDTA for 10 min in RT and the cell suspension was further diluted with 5ml PBS before separating cells by centrifugation and dissociating them with 0.1ml Accutase enzyme for 5 minutes (due to aggregation of cells). A sufficient amount of media was then added, and cells were carefully re-suspended and counted by Hemocytometer.

## 3 Results

The first aim was to generate mesodermal progenitor cells from human induced pluripotent stem cells (hiPSCs) [16]. After characterization of hiPSCs by detection of pluripotency markers Oct4 and Sox2 (Fig.1 A1-A3) using immunofluorescence staining (IF), the conversion of these cells into mesodermal progenitor cells (hiMPCs) was performed under directed differentiation conditions according to a protocol previously established in our lab [16]. After 3 days of culturing in mesodermal induction media, the cells were dissociated using Accutase and IF detecting mesodermal progenitor cell markers such as vascular endothelial growth factor receptor 2 and Brachyury were performed (Fig.1 B1-B3).

We then studied the behavior of mesodermal progenitor cells (hiMPCs) after formulation into alginate-based bioinks and subsequent extrusion by a pressure-assisted dispersion technique through a nozzle system. Initial viability assessment demonstrated that cells survive this procedure at best within a sodium alginate+collagen type1 bioink (Fig. 2C). After extrusion of the bioink containing hiMPCs into the silicone molds, the resulting constructs were cultivated in endothelial cell differentiation medium. In order to promote the differentiation and to increase the number of vascular cells, particularly of ECs, further factors such as L-Glut, Vitamin C, Penstrep, Rock inhibitor and particularly VEGF were added to the cell culture medium. The added factors are also beneficial for the induction, promotion and maintenance of vascular morphogenesis including capillary-like tube formation [25, 26].

**Figure 2:**
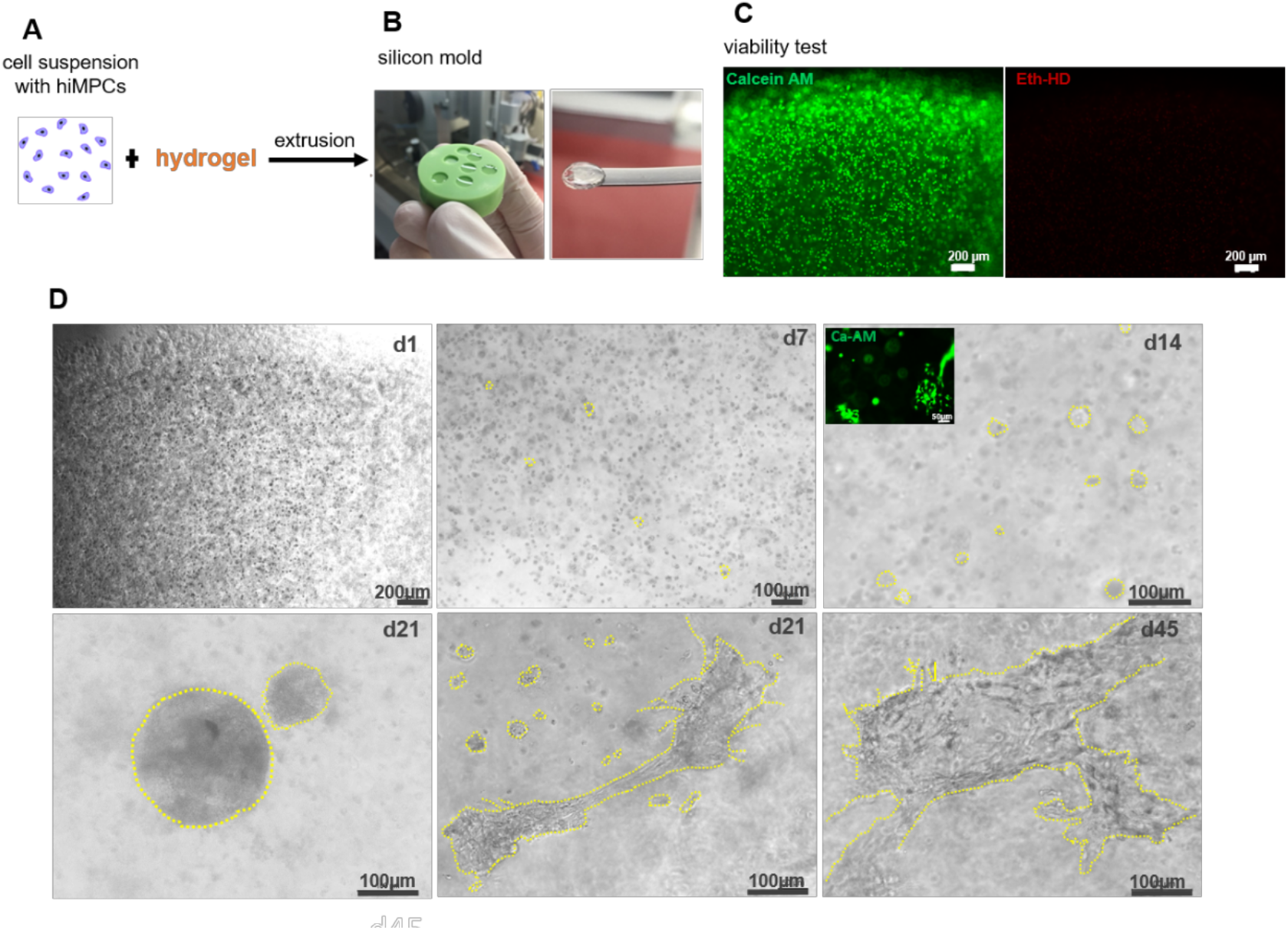
Extrusion-based dispensing of bioinks containing hiMPCs into silicone molds, initial viability test and phase contract microscopic study of cultured constructs **A**. Cell suspension of hiMPCs formulated into alginate alone or alginate + collagen type I as bioink, **B**. Disc form after extrusion into the silicon molds **C**. Initial viability testing by application of Calcein-AM and EthD-1. **D**. Phase contrast microscopy of cultured constructs at different culture time points as indicated. Morphogenetic events with single cells at day 1, small cell spheres at day 7, large spheres and cell cords at day 14, different sized spheres and cell cords with branching at day 21 and large branching cell cords at day 45.

3D bioprinting techniques need bioinks that are biocompatible, non-immunogenic and non-toxic, and further require cells that have the capacity to deliver tissue- and/or organ-specific parenchymal cells in order to be able to enter tissue and organ morphogenesis. Alginate as a frequently used bioink attracts a lot of attention and is preferred by many groups not only for the features listed above, but also due to its properties such as easy handling, fast cross-linking, controllable biodegradation as well as easy producibility [27]. The physical and chemical properties of the hydrogel should allow and support basic cellular behavior such as attachment to the matrix as well as cellular proliferation and differentiation. For initial testing of the extrusion process sodium alginate alone was chosen as bioink. Cells and biomaterials were mixed carefully, free from air bubbles and placed into printing cartilage. The cell-loaded bioinks were extruded into disc-shaped silicon molds. During extrusion, cells are subjected to different shear stress within the nozzle. To reduce cell stress, it was necessary to determine the optimal nozzle size, suitable extrusion parameters and most favorable cell density to preserve cell vitality and basic cellular behavior within the hydrogel after extrusion. To perform the extrusion, a homogenous, sterile 2% alginate hydrogel was prepared. Different suspensions of hiMPCs (2.5*10^6^, 5*10^6^ and 9*10^6^cells/ml) were mixed with the hydrogel. The cell-loaded gel was extruded through the nozzle under a continuous pressure of 100 kPa. The gel discs were then cross-linked and transferred into cell culture flasks for further culturing as visualized in figure 2B. Based on results we obtained from such optimization analyses, we decided to extrude the cells through 25G ¼ inch lock tip nozzle that preserved at the best the cellular viability and functionality.

Generally, extrusion of more than 5*10^6^ cells/ml resulted in a cell survival time longer than 3 months within the alginate gel while morphogenetic events were limited. Extrusion of less than 2.5 million cells/ml into alginate-based hydrogel alone resulted in a significant reduction of cellular survival time below 2 weeks. This indicates that a sufficient cellular density is needed in order to probably achieve a sufficient cell-cell communication within the hydrogel either by direct cell-cell interaction and/or in a paracrine manner via secreted factors from differentiating cells, e.g. ECs, pericytes or SMCs that might support the morphogenetic events and thus, also cell survival. In cases of using alginate+collagen type I as bioink as mainly applied in the studies here, a cell density ranging between 2.5*10^6^ to 10*10^6^ cells/ml turned out to be the best option.

Then, we studied morphogenetic events under phase contrast microscopy. We observed the formation of 3D cell spheres in both alginate-based bioink alone and alginate+collagen type I bioink starting around culture day 5-7. However, a network of elongated and enlarged cell cords was only observed in alginate+collagen type I hydrogel (Fig. 2D and 3A). It is well described that *de novo* development of blood vessels during the embryogenesis starts also with formation spherical cell aggregates by specialized mesodermal progenitors that form, the so-called “blood islands”, that serve as the source for both endothelial and hematopoietic cells [28, 29].

As a biomaterial, alginate turned out to be suitable for keeping hiMPCs viable. However, the gel is mechanically tough and not adhesive enough to provide cell attachment. For that reason, it did not support cell migration, a process that is essential for initiating vascular morphogenesis. To this end, we decided to modify the alginate bioink by blending with collagen type 1 (col type 1) to support the cellular migration, proliferation and thus, morphogenetic events. Indeed, phase contrast microscopic observation at culture days 7-9 after extrusion revealed elongated cell cord formation and morphogenetic processes including the formation of a vessel-like branching pattern. These data suggest that the hiMPCs actually require the presence of collagen type 1 in order to migrate, proliferate, interact and form 3D structures within the alginate-based hydrogel. hiMPC containing collagen type 1 gelation should be dispensed into alginate as fast as possible in order to avoid non-homogenous dispersion of the bioink. This is critical parameter that should be taken into account while preparing cell-loaded gel prior to extrusion.

The viability analyses using Calcein AM at three incremental time points: 1, 3, and 7 days allowed beside the cell survival assessment also to follow some morphogenetic events such as the formation of cells spheres, cell cords and their branching pattern and the formation of hollow and tubular structures (Fig. 3A-C).

**Figure 3:**
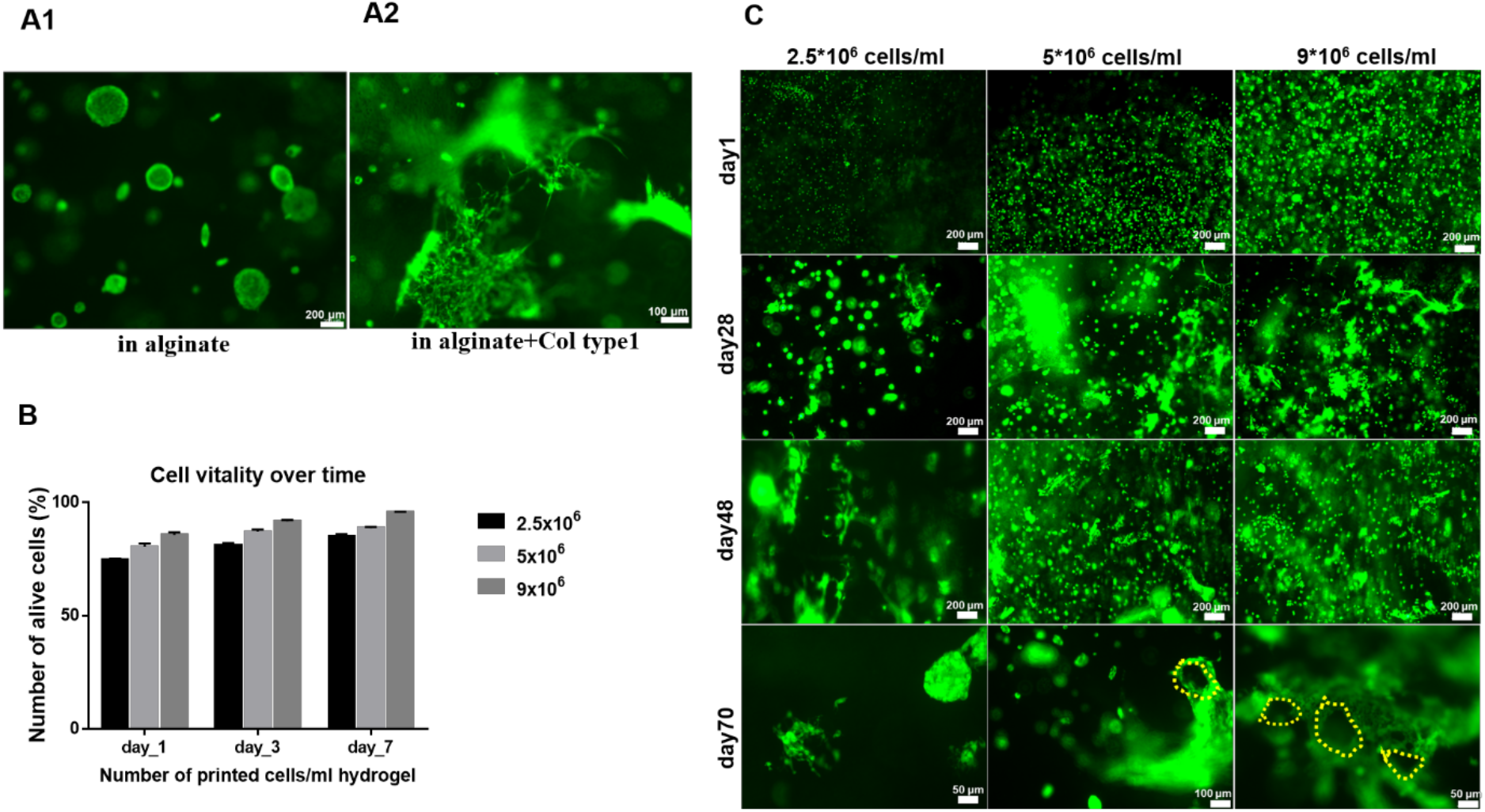
**A**. Calcein-AM-fluorescent cell viability images of the extruded cells (A1) in alginate at day 100 and (A2) in alginate+col type-I at day 70. **B**. Percentage of live cell counts of the extruded cells within alginate+collagen type1 hydrogel at culture day 1, 3 and 7 (bar colors indicate the cell density). **C**. Fluorescent cell viability images of the extruded hiMPCs at different density within alginate+collagen type I hydrogel. The extrusion parameters: 25G (ID = 340 µm) ¼” nozzle, one bar, RT, into silicone mold. Yellow dotted lines mark hollow structures formed by hiMPCs that are already visible after Calcein-AM fluorescent labelling.

These results clearly suggest that the enrichment of alginate with collagen type I results in a cell responsive bioink that supports cell attachment, cell migration and cell-cell as well as cell-gel interaction better than alginate-based hydrogel alone. Moreover, the elongated cords displayed network formation to some extent (Fig. 3C). To this end, almost all basic steps that are needed for vascular morphogenesis and vessel network formation were achieved using alginate plus collagen type I as bioink. Furthermore, the optimal extruded cell density reduced the adverse effect on cell viability and survival and resulted in an improvement of the aforementioned parameters in a ratio up to >75% (Figure 3B-C).

In order to better study the mentioned morphogenetic events, the printed constructs were embedded in paraffine and cross sections (10 µm) of these tissue blocks were used for H&E staining. Light microscopic analyses revealed time-dependent morphogenetic changes. The cultures started with single cells distributed randomly within the gel at culture day 1, followed by the formation of cell spheroids at day 10 and a branching vascular-like channel system at day 25 after bioprinting (Fig. 4A). These morphogenetic steps clearly suggest that hiMPCs recapitulate a differentiation program that is well known from the *de novo* embryonic vascular development by vasculogenesis as described previously [30]. Remarkably, lumen formation within cell spheroids can be detected already at culture days 7-10 (Fig. 4A). These events are progressing with further culture time and result in a) stabilization of the wall of the vessel-like channels, e.g. by assembly of peri-endothelial cells into the wall and b) branching, probably due to angiogenic sprouting of new vessels from the already formed ones as visualized by H&E staining at day 14 (Fig. 4A). The flattened and elongated nuclei of cells lining the luminal surface indicate their endothelial cell-like identity. The H&E stained inset in figure 4A that was captured from a section of day 21 after extrusion clearly displays lumen formation and a multilayered vessel-like wall structure (Fig. 4A). Furthermore, the paraffine sections were also stained with Picrosirius red in order to visualize the collagen matrix (Fig. 4B). This staining revealed a dense deposition of collagen around the vascular-like branching network that also enabled the recognition of lumen formation in some areas of this network. We assume that this densely deposited collagen reflects not only the collagen type I that was added to hydrogel but also collagen components that were produced by the cells themselves while forming vessel-like structures.

**Figure 4:**
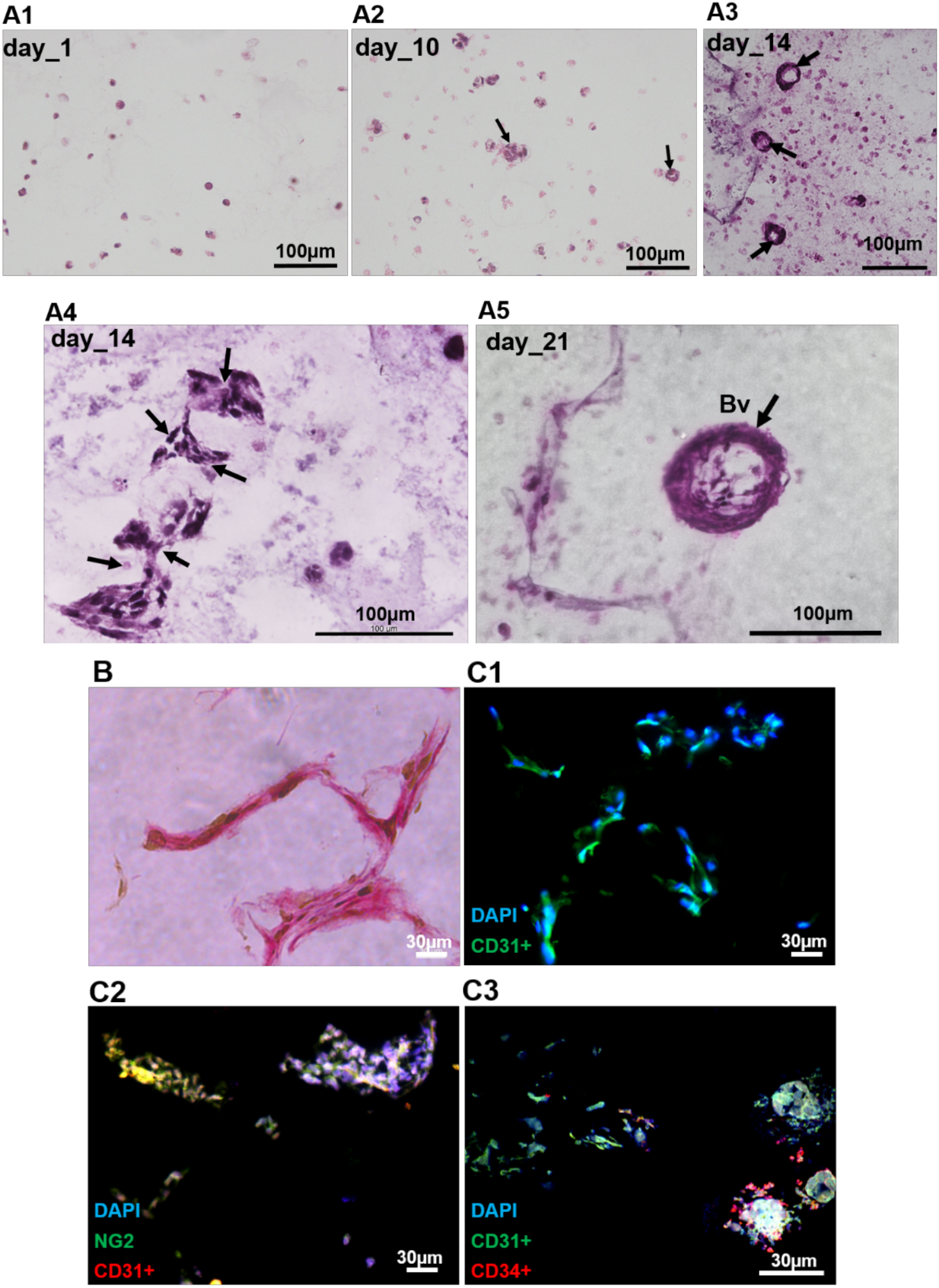
**A**. H&E staining of extruded constructs of alginate+collagen type I containing hiMPCs at day 1, 10, 14 and 21 of culture. Single cells at day 1 (**A1**), spheroid formation at day 10 (arrows, **A2**), formation of vessel-like structures (arrows, **A3**), connected and branching small vessels (arrows, **A4**), relatively large vessel with multilayered wall structure (Bv, **A5). B**. Picrosirius staining of the extruded constructs after 2 weeks demonstrating vessel-like network formation. The image was taken with 40x magnification. **C**. Immunofluorescence staining for CD31, CD34 and NG2 on sections of the extruded constructs of alginate+collagen type I containing hiMPCs.

During blood vessel formation, endothelial progenitor cells called angioblasts emerge as scattered cells in the mesoderm and aggregate to form cords that predetermine future blood vessels. The endothelial cells that are derived from these progenitors reorganize into blood-carrying tubes forming a primary vascular plexus by a process that is termed vasculogenesis. Subsequently, this initial primary vascular network is remodeled and extended by mechanisms that govern the further vascular maturation up to formation of a hierarchically organized vascular system [30-32]. VEGF signaling through the VEGF receptors-1 and -2 is the major mechanism that regulates almost all aforementioned steps of vascular formation and remodeling in health and diseases [33]. Thus, we added VEGF in all stages of culturing the extruded constructs containing hiMPCs. In addition to increase the cell–cell interactions which is required throughout the formation of the vasculature and must occur in a precise spatio-temporal sequence [34], the constructs were cultured in endothelial growth medium. Moreover, signaling of non-endothelial cells, e.g. peri-endothelial cells in a close vicinity to the ECs as well as signals from the extracellular matrix are crucial for maintenance and further maturation of newly created blood vessels [35, 36]. Our data clearly suggest that such processes, mechanisms of cell-cell- and cell-matrix-interaction are not only essential in *in vivo* vascular formation but also indispensable for vascular bioprinting.

The basic indispensable step of vascular maturation is the tight integration of peri-endothelial cells such as pericytes and smooth muscle cells into the vessel wall [25]. Next, we wanted to improve this step, in order to achieve a multilayered vessel wall and thus varied the number of cells being extruded in bioprinting. These analyses revealed that increasing the density of hiMPCs, e.g. up to 9*10^6^ cells/ml resulted in formation of vessel-like channels of different size. A part of these channels displayed a multilayered wall structure. Of note, extrusion of hiMPCs at a density of 2.5*10^6^cells/ml resulted in capillary-like micro vessels missing a multilayered wall structure and the vessel formation took place after a significantly longer time period (almost 2 months). These data underline again the impact of using an optimized cell density in bioprinting of blood vessels.

Next, we wanted to characterize the cells forming the vascular-like channels at the immunophenotypic level using immunostaining on paraffine sections for markers that are specific for endothelial cells such CD31, for peri-endothelial cells such as NG2 and αSMA, and for vascular progenitor cells such as CD34 [13, 14, 37, 38], The presented results demonstrate that indeed all vascular cell types are present in the wall of the vascular-like channels developed after extrusion of hiMPCs into the alginate+collagen type I matrix (Fig. 4C). Moreover, the spatial organization of these cell types within the wall of vessel-like channels was very similar to that *in vivo* as CD31^+^ ECs lined the luminal surface of the vascular channels, the NG2^+^ or aSMA^+^ cells were found to cover the endothelial layer from outside and more surprisingly a part of CD34+ cells was found in the outermost layer of the vessel wall, again similar to the adventitial layer of blood vessels *in situ* (Fig. 4C3).

Based on the aforementioned results from histological and immunohistological studies, we aimed at analyzing these vascular channels in more detail at the ultrastructural level using electron microscopic imaging via TEM and serial block face REM. These analyses confirmed entirely our histological observation suggesting the presence of whole spectrum of vessel types representing of different parts of the vascular hierarchy such as capillaries with and without pericyte coverage and basement membrane as well as large and mid-sized vessel-like channels that displayed a multilayered vessel wall (Fig. 5). Already in toluidin blue-stained semi-thin sections of our constructs revealed capillary-like structures exhibiting cells lining the lumen (intima) and a cell that covered this endothelial lining from the outside (Fig. 5A). Moreover, the morphological appearance of these covering cells is similar to the pericytes *in situ* (Fig. 5A and 5C1). As also visualized in the same figure panel, there are also capillaries that are constructed by ECs only (Fig. 5A, B2 and B3). Since we observed such structures in almost all tissue blocks, we assume that the vessel formation is taking place in relatively large area within the alginate+collagen type I gel. The subsequently performed electron microscopic analyses confirmed these observations at the ultrastructural level (Fig. 5A-5C). Some capillary-like channels displayed less flattened cells indicating that they are structurally still at an immature state (Fig. 5B). In order to visualize a more mature capillary-like vessel in its whole circumference at relatively high magnification, the single TEM figures were fused and the entire endothelial lining of the capillary lumen is demonstrated (Fig. 5C). Higher magnification revealed that the EC layer is anchored at a basal lamina-like condensed matrix that is tightly attached to the basal side of the intimal layer (Fig. 5C3). Moreover, confirming our histological analyses on paraffin sections, also TEM studies revealed that a part of blood vessels displays a multilayered wall structure (Fig. 5C). We could identify at least three concentric layers: the innermost intima that is followed by media and the outermost adventitia (Fig. 5C3). Furthermore, the section morphology of the cells in these layers clearly visualize that ECs are more oriented in the long axis of the channel, the media cells are oriented circularly and the adventitia-like cells display many thin processes that are embedded in a collagen matrix that is apparently structured by the vascular wall cells (Fig. 5C3).

**Figure 5:**
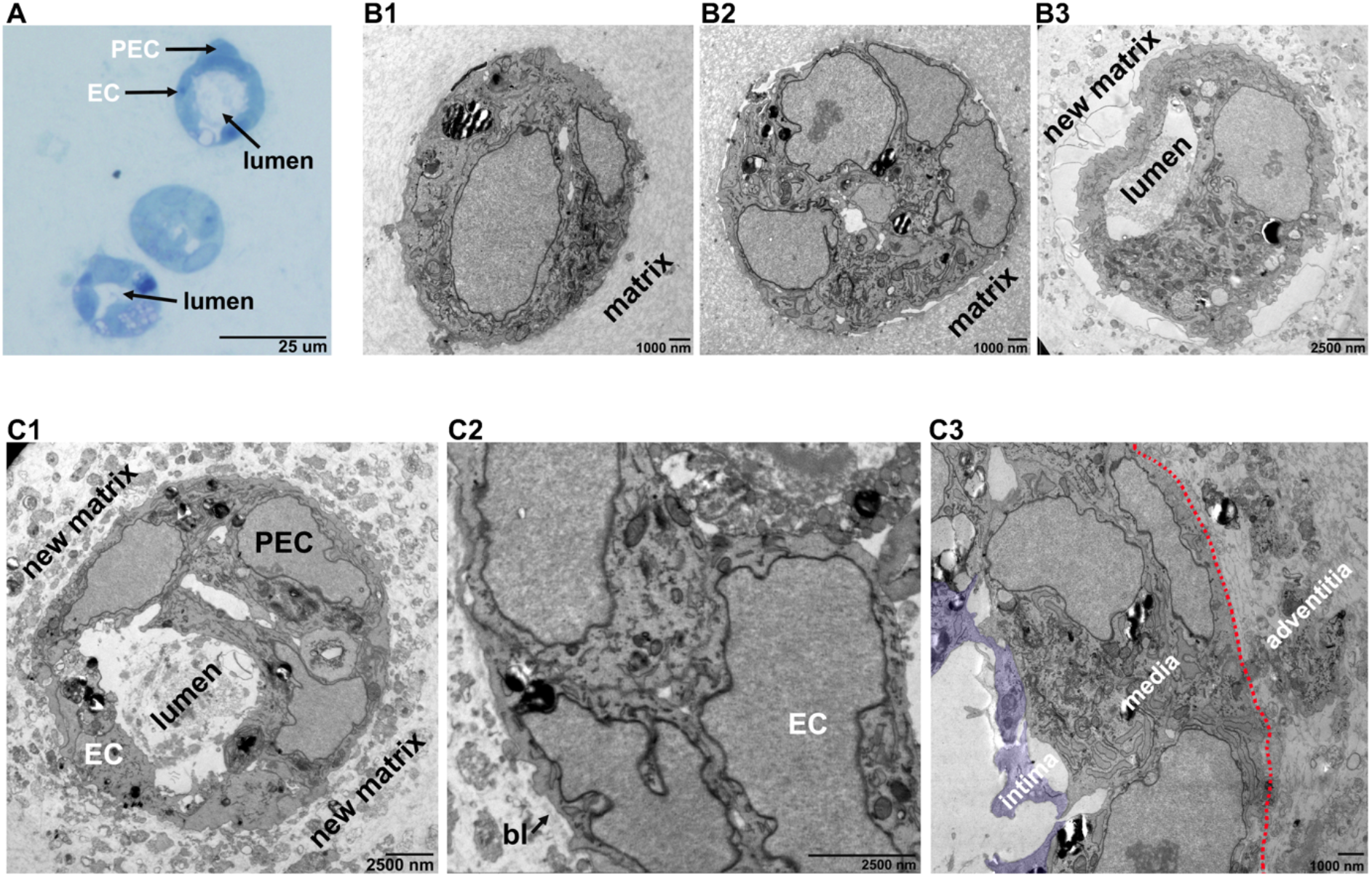
Semi-thin sectional and ultrastructural evaluation vessels generated by extrusion- based shape-free printing of hiMPCs+alginate+collagen type I into silicone mold. **A**. Semi-thin section analyses results after *<u>toluidin blue</u>* staining (1 µm sections) display vessel-like tubes lined by ECs and PEC (peri-endothelial cell, e.g. pericyte). **B**. Electron microscopic analyses show single cells (B1), cell spheroid (B2) and a capillary that consist of only a single layer of ECs forming a lumen and new extracellular matrix surrounding this capillary (B3). **C**. Electron microscopic studies show vessels with ECs and PECs. Capillary-like vessel with ECs and PEC and new matrix deposition around these vessels (C1). Blood vessel with single innermost layer of ECs that is covered by single PEC form outside and that in turn is underlined by a dens basal lamina as indicated (arrow and **bl**) (C2), Blood vessels innermost intima (marked by purple color), two-three concentric cell layers of media and the outermost adventitia as separated from the media by the dotted red line (C3).

In order to better visualize the vessel-like lumen formation we used serial block face electron microscopy (SBF-SEM) and performed 3D reconstruction about a distance of 7.5 µm (Fig. 6). These analyses revealed that hiMPC-derived vascular cells formed complex lumen with branching pattern that is provided by thin cellular processes which subdivide a lumen into two further lumen and by this, create a complex communicating channel system. While the further maturation of these channels depends on perfusion and further vascular maturation, these data demonstrate morphogenesis of vessel-like lumen and their branching, e.g. similar to the vessel branching processes *in situ* [39, 40].

**Figure 6:**
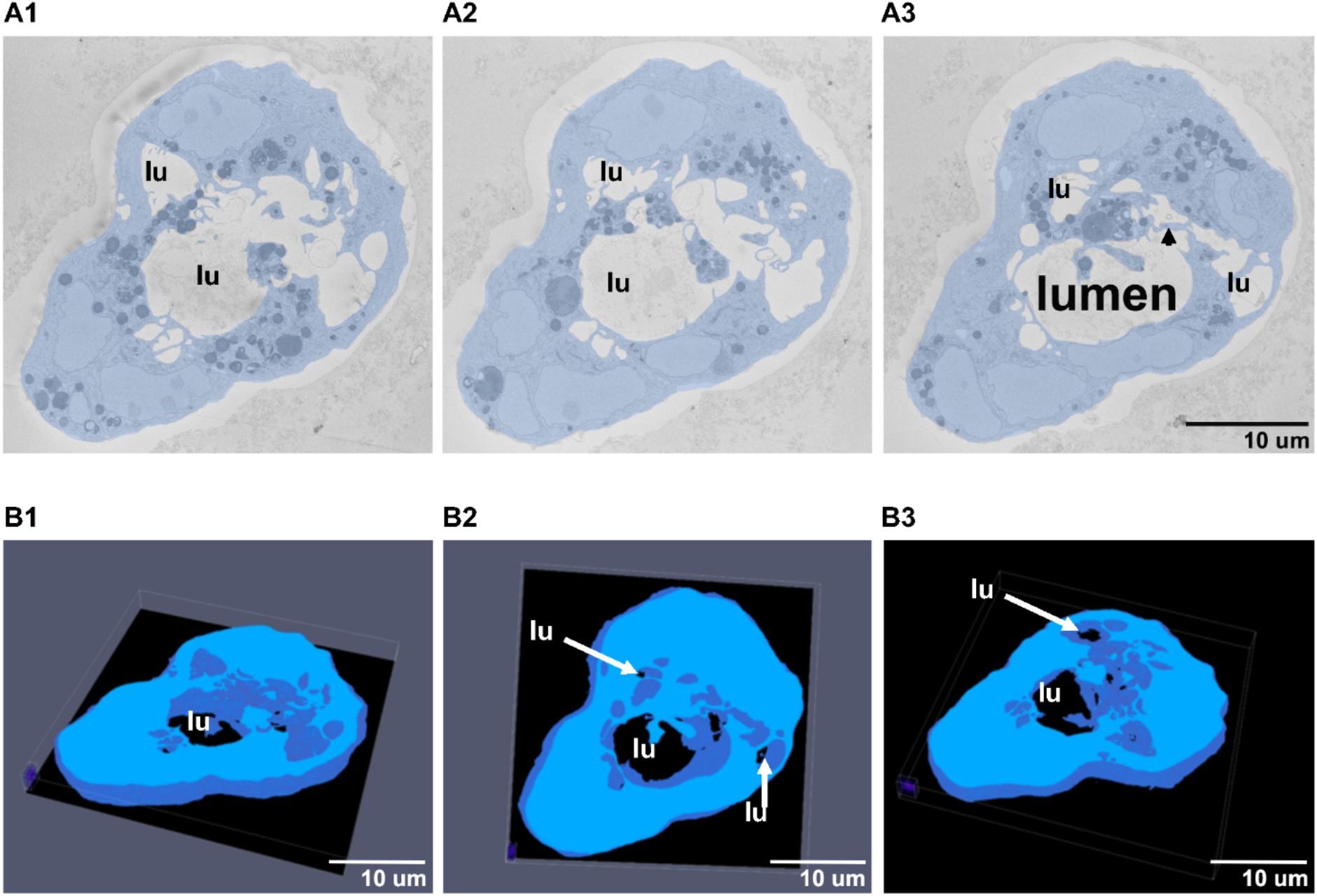
SBF-SEM imaging of branching lumen (lu) formation by hiMPC-derived vascular cells after extrusion into silicone molds. **A1-3**. Selected images from single SBF-SEM slices showing hiMPC-derived endothelial-like cells (blue) that form a complex lumen (**lu**) with thin cellular processes leading to lumen branching (arrowhead, **A3**). **B1-3**. Large-volume reconstruction of the SBF-SEM slices presented in (A) showing the branching pattern by thin processes of endothelial-like cells that separate the lumen (**lu**) from each other. SBF-SEM dataset: Reconstruction of a 7.5 µm volume: 151 sections, 10nm x 10nm x 50nm voxel size. Source data are available for this figure. Scale bars as indicated.

## 4 Discussion

The present data demonstrate an artificial generation of a hierarchically organized vascular system after extrusion by combining vascular stem cell technology with bioprinting technology that allowed to mimic the basic steps of *de novo* vessel formation during embryogenesis. Briefly, we demonstrate, as also summarized in the graphical representation (Fig. 6), that a) hiPSC-derived mesodermal cells (hiMPCs), that were recently shown to have a vasculogenic potential [16] were formulated into hydrogels composed by alginate alone or alginate+collagen type I and survived sub-sequent the extrusion-based dispensing into a silicon mold, b) 7 days after extrusion hiMPCs differentiated and underwent morphogenetic events such as formation of 3D spheroids in several places within the hydrogel, c) few days after spheroid formation, elongated cell cords emerged that displayed also morphological features of branching and network formation, e.g. connecting neighboring spheroids, d) at culture days 15-21 the 3D cell cords were enlarged displaying more complex branching and networking, e) conventional histological H&E staining on paraffin sections of the aforementioned constructs revealed vessel-like channels, f) immunostaining of the sections for the endothelial marker CD31 and CD34 as well as for NG2 and aSMA as pericyte and smooth muscle cell marker confirmed the vascular nature of the channels and the right spatial organization of endothelial and peri-endothelial cells in their wall, and finally g) electron microscopic studies of these constructs revealed capillaries, small vessels with single peri-endothelial cells and larger vessels with three-layered wall structure such as intima-, media- and adventitia-like layers. Furthermore, SBF-SEM analyses with segmentation and 3D reconstruction revealed vessel-like lumen formation that is subdivided into two further lumens by cellular processes which create a communicating vessel-like channel system.

The present data suggest that using vascular progenitors that have the capacity to deliver all mature vascular wall cell types such as endothelial cells, pericytes, smooth muscle cells and mesenchymal cells or fibroblasts in biofabrication of blood vessels is a promising approach to generate artificial vascular system in bioprinted tissues and organs. At the biological level, this approach uses a) the vasculogenic potential of the vascular progenitor cells instead of using mature vascular cells, and b) mimics the embryogenesis of blood vessels by combining the aforementioned cellular potential with material and the techniques of biofabrication in order to achieve not only a rigid pipe but rather a plastic and adaptable vessel system.

Tissue vascularization is an indispensable prerequisite for proper development and function of tissue as well as organ. The primary hypothesis of the present work was to test the capacity of vascular progenitor cells to form vessel-like structures by self-assembly after being formulated in bioink, e.g. alginate+collagen type I and subsequent extrusion into a silicon mold as performed here. We chose this setup in order to unambiguously explore whether the hiMPCs do the expected job after being exposed to sheer stress through syringe and within a matrix that is different as the matrix *in situ*. Furthermore, our second aim was to test to which extend the hiMPCs can enter into morphogenetic events within the hydrogel, there needed for formation of vessel-like structures. Moreover, the best desired outcome would be the formation of blood vessels that exhibit 1) different size, e.g. capillaries and larger vessels, 2) a multilayered well structure and 3) a vascular network that is organized in hierarchic manner as blood vessels *in situ*. To our best expectation, all aforementioned goals were met by using hiMPCs seeded into alginate+collagen type I and extruded shape-free into silicon mold. As documented in the main body of published work regarding biofabrication of blood vessels, mature vascular cells such as EC, e.g. HUVECs [41, 42], human microvascular endothelial cells (HMVECs) [43] or smooth muscle cells [44] as well as ECs plus fibroblasts [45], HUVECs plus mesenchymal cells plus fibroblasts [46] and HUVECs plus smooth muscle cells [8]. While these cell types have a vascular phenotype and thus, can be expected to form easily vascular structures, the mature state or character of these cells has to be considered when using them for biofabrication of new vessels. Due to their mature state ECs, SMCs or fibroblasts cannot run the basic molecular and cellular program as e.g. the so-called angioblasts of mesodermal origin do during *de novo* embryonic vascular development by vasculogenesis [47]. It has also to be considered that the developmental potential of the mesodermal angioblasts is activated and controlled by essential growth factors and cytokines like vascular endothelial growth factor (VEGF), basic fibroblastic factor (bFGF), platelet-derived growth factor (PDGF) and angiopoietin-Tie-2 signaling. Among these factors, the VEGF-VEGF-R2 (Flk1) signaling system is the master regulator of vasculogenesis as studies on gene knock-out models revealed that even mice with a heterozygous deletion of only one allele of VEGF gene was embryonically lethal due to the severe vascular defects on very early stage of embryonic development [48, 49]. VEGF is also needed not for initial vascular formation but together with PDGF and angiopoietins it also needed for further vascular maturation, e.g. by assembly of peri-endothelial cells such as pericytes and smooth muscle cells into the wall of new vessels which lead to vascular stabilization and survival [50]. Based on aforementioned knowledges, we brought not Flk1-postive mesodermal progenitors and bioinks together in a formulation and further added VEGF to our constructs after extrusion and under culture conditions, in order to initiate the basic program of hiMPCs to formation of blood vessel like structures. Once initiated, mesodermal cells give rise to generation of all vascular wall cell types such ECs, SMCs and MSCs/fibroblasts that in turn can interact with each other and orchestrate the further steps of vascular morphogenesis including a hierarchic vessel system containing both micro and macrovessels with multilayered vessel wall as demonstrated in this work. We recently used hiMPCs in organoid models where they also provided vascularization [16, 51] underlining the basic vasculogenic potential of hiMPCs. Remarkably, in comparison to the organoids where we observed vessels with only one layer of peri-endothelial cells we achieved in the current study the formation of vessels with multilayered tunica media as demonstrated by histology and electron microscopic analyses. Moreover, in vessels with a diameter 100 – 200 µm or above, we also observed an adventitia-like outermost layer similar to the vessels *in situ*. The CD34^+^ immunophenotype of a part of these adventitial cells underlines also their spatial organization within the vessel wall like blood vessel *in situ* and the CD34^+^ vascular adventitial cells have been shown to have angiogenic and vasculogenic potential even in adult vessels [13, 52]. The advantage of this approach, as we initially postulated, lies in the fact, that we indeed could mimic the basic steps of embryonic vasculogenesis in a shape-free biofabrication procedure of creating artificial blood vessels as graphically visualized below (Fig. 6). With this, our approach opens a new avenue for biofabrication of blood vessels by combining technologies of shaped biofabrication+vascular progenitors (iPCs-derived or primary vascular progenitors) +vasculogenic factors. This would enable to improve the simultaneous vascularization of biofabricated tissues and organs on the one side but also improve the cardiovascular as an organ model itself.

**Figure 6:**
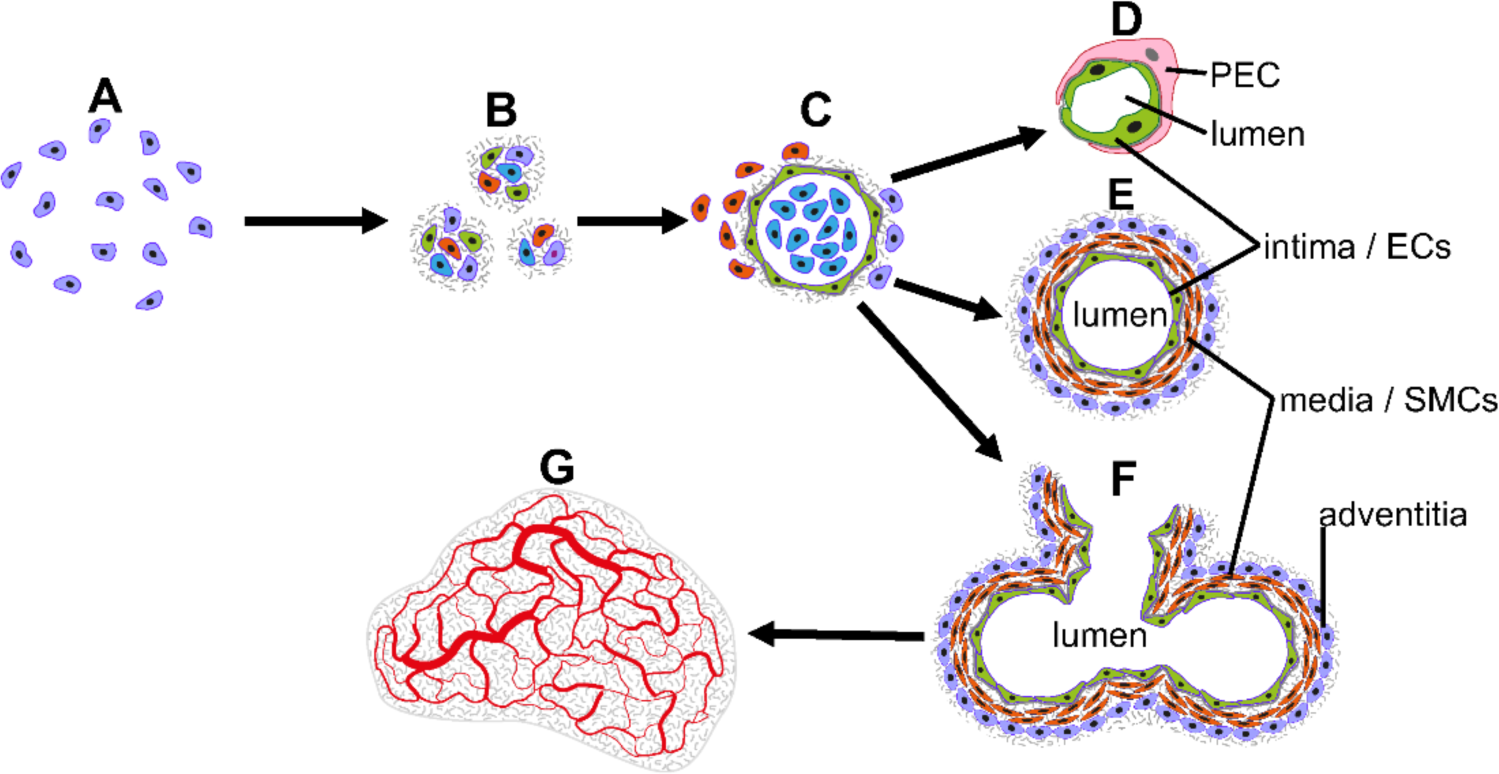
Graphical representation of vascular morphogenesis steps after extrusion-based shape-free printing of hiMPCs that were seeded into the alginate+collagen type I hydrogel into a silicone mold. hiMPCs (blue) seeded into the hydrogel (**A**), cell sphere formation (like embryonic angioblasts) within the hydrogel (**B**), lumen formation by relatively flattened cells like ECs (green) with some surrounding cells that probably committed to peri-endothelial cells (red) (**C**), differentiation and further maturation of vessel-like channels to a capillary phenotype (**D**) and, branching vessels with multilayered wall structure exhibiting an intima, media and adventitia (**F)** Within the silicone mold, this vessel-like channels formed a 3D network composed by small and large vessels that displayed a hierarchic organization schematically visualized (**G**).

## 5 Conclusion

With the current study we introduce a new approach in the field of blood vessel biofabrication using a) human iPSC-derived hiMPCs which were formulated in alginate+collagen type I bioinks and printed into molds by extrusion, and b) culture of these constructs in endothelial growth medium by adding VEGF in order to induce the vascular potential of hiMPCs mimicking the *de novo* blood vessel formation during embryogenesis. We demonstrate that this approach resulted in formation of small and large vessels with multilayered wall structure displaying intima-, media- and adventitia-like layers. Moreover, the vessels were organized in a hierarchic vascular network by branching in a similar pattern as the vascular system *in situ*. Adapting this approach to other types of bioinks as well as techniques of bioprinting allowing shaped biofabrication of blood vessels in tissue- and organ-specific pattern under adding essential factors that are indispensable for proper vascular morphogenesis such VEGF, PDGF and angiopoietins in future studies would enable a major step forward in achieving the goal of vascularization of biofabricated tissues and organs.

## Acknowledgement

This research was funded by the Deutsche Forschungsgemeinschaft (DFG, German Research Foundation) – Project number 326998133 – TRR 225 (subproject B04)

